# *BRCA1* is a molecular correlate of cell proliferation in human brain development and in Group 3 and 4 medulloblastoma

**DOI:** 10.1101/2025.04.03.646803

**Authors:** Ian Cheong, Xinghan Sun, Leo Lau, Nishka Kishore, Andrea Senff-Ribeiro, Ander Diaz-Navarro, Hana Hajari, Ellen Mak, Rhea Ahluwalia, Luca Bianchini, Jaskirat Singh Sandhu, Michael D. Taylor, Lena Kutscher, Lincoln D. Stein, Shraddha Pai

## Abstract

The role of the BRCA1-mediated DNA damage repair pathway in regulating human brain development remains unknown, although it has been studied in mouse development. We report evidence for breast cancer type 1 susceptibility protein (BRCA1) being a molecular correlate of proliferation in human neural progenitor cells and in medulloblastoma (MB), a malignant pediatric hindbrain cancer whose cells resemble undifferentiated neural stem cells. In a computational search for molecules potentially keeping Group 3 (G3) and Group 4 (G4) MB tumour cells in a state of stalled differentiation, BRCA1 emerged as a leading candidate gene. We surveyed four independent transcriptomic datasets collectively spanning 142 human developing brain samples, multiple brain regions, and over 1.7 million single cells, and found that BRCA1 transcription is consistently enriched in human neural stem and progenitor cells, relative to differentiating or mature neurons. Across the human lifespan, BRCA1 expression is enriched in the brain during early development, particularly the first trimester of gestation. By analyzing 714 tumours, the largest transcriptomic survey of MB tumours to date, we found that BRCA1 expression is increased in carriers of isochromosome 17q (i17q) aberrations and in G4 MB tumours. Increased BRCA1 expression is associated with worse prognosis in G3 and G4 MB. In the developing cerebellum as well as in the cancer context, BRCA1 expression is correlated with transcription of the cell cycle and DNA damage repair pathways. When considered with previous mouse studies, our work is consistent with a model in which BRCA1 promotes genome surveillance in neural progenitors during human brain development and in G3 and G4 MB tumour growth, thus supporting proliferation.

## Introduction

A hallmark feature of pediatric brain tumours is the molecular similarity of tumour cells to undifferentiated neural stem and progenitor cells normally present in early neurodevelopment^1–6^. This observation made across multiple forebrain and hindbrain tumours, such as diffuse intrinsic pontine glioma, medulloblastoma, ependymoma, and atypical teratoid rhabdoid tumours, has led to the working model that a common mechanism in the oncogenesis of pediatric brain tumours is stalled differentiation in development. Delineating neurodevelopmental mechanisms and their dysregulation in cancer will advance knowledge of molecular control of human neurodevelopment, improve patient stratification, and suggest pathways for rational therapy design.

Stalled differentiation is a leading hypothesis for the origins of medulloblastoma. Medulloblastoma is the most common malignant pediatric brain cancer and lacks targeted molecular therapies. The current standard of care, which consists of surgical resection followed by chemotherapy or craniospinal radiation, leaves survivors with a diminished quality of life^7,8^. Over 65% of medulloblastomas fall into two broad, overlapping molecular subgroups, Group 3 and Group 4, that are poorly understood at a molecular level and have the worst outcome. Group 3 and Group 4 medulloblastoma tumours resemble neural progenitor cells of the rhombic lip of the developing cerebellum, and a leading hypothesis in the field is that these tumour cells originate from stalled differentiation of the rhombic lip^1,2^. This finding has promising therapeutic potential, as activation of cerebellar neuronal differentiation pathways results in upregulation of differentiation markers in tumour cell lines and in mouse models of medulloblastoma, and reversal of tumour growth^2,9^. However, the full set of developmental pathways aberrantly upregulated in Group 3 and 4 medulloblastoma tumours — which comprise 60% of all cases — remains unknown. Separately, human cerebellar development remains poorly studied, as most molecular studies to date have been performed in the mouse. By uncovering the developmental origins of medulloblastoma, we aim to better understand the dysregulated pathways underlying stalled differentiation while also helping to advance knowledge of human cerebellar development.

Here we provide evidence for BRCA1 as a supporter of proliferation in human neural stem and progenitor cells and in Group 3 and 4 medulloblastoma tumour cells. We report widespread expression of BRCA1 in human neural stem and progenitor cells of the forebrain and hindbrain, and highest expression during the first trimester of gestation. Expression of BRCA1 is correlated with cell cycle and DNA damage repair pathways, consistent with its established role in these processes in adult tissues (see Roy *et al.*^10^ for a review). In medulloblastoma, we find that BRCA1 expression is particularly enriched in carriers of the i17q chromosomal aberration and in Group 4 tumours, and that increased BRCA1 expression is associated with worse prognosis in Group 3 and 4 medulloblastoma tumours. Collectively, our results are consistent with a model wherein BRCA1 acts as part of the DNA damage repair pathway to protect genome integrity during the rapid cell proliferation that accompanies the early growth of the human brain.

## Methods

### Computational pipeline to predict candidates for stalled differentiation

Processed rhombic lip (RL) cells and cell cluster annotation from the Aldinger scRNA-seq dataset of microdissected developing human fetal hindbrain^11^ were obtained from Dr. Michael D. Taylor.^2^. Seurat 4.3.0.1 was used to compute genes differentially expressed between the “rhombic lip subventricular zone” (RL-SVZ) and “early unipolar brush cell” (early UBC) cluster and adjusted for sample sex (parameters were set as follows: *assay = “SCT”, test.use=“LR”, latent.vars=“sex”, recorrect_umi = FALSE*). Genes with an adjusted p-value < 0.05 and log2 fold-change > 0 were deemed upregulated in RL-SVZ. Transcription factor binding sites enriched in RL-SVZ upregulated genes were identified using AME (MEME v5.4.1)^12^ with the HOCOMOCO v11 Core Human motif database (HOCOMOCOv11_core_HUMAN_mono_meme_format.meme). The list of protein epigenetic factors was downloaded from the EpiFactors database^13,14^ v2.0 (URL: https://epifactors.autosome.org/public_data/v2.0.zip). The list of human transcription factors was obtained from Lambert *et al.*^15^ (URL: https://humantfs.ccbr.utoronto.ca/download.php). The set of genes mutated or overexpressed in medulloblastoma were taken from Supplementary Table 5 of the Hendrikse *et al.* paper ^2^; genes with Q < 0.05 for OncoDrive FML or MutSig, for either Group 3 or 4 MB tumours were included as potential MB drivers. The full list of genes is included in Supplementary Table 17. Gene expression and CRISPR impact on proliferation on the D425, a human Group 3 MB cell line, was downloaded from DepMap^16,17^.

### Developmental scRNA-seq analysis

Braun *et al.* (2023) human first trimester whole brain: Processed scRNAseq expression data for the human first trimester brain was obtained from Braun *et al*. (2023) (over 1.6 million cells, https://github.com/linnarsson-lab/developing-human-brain?tab=readme-ov-file). The R package *rhdf5* was used to analyze the data. Cell type labels were used as assigned by Braun *et al*. Known markers to characterize different lineages such as radial glia (marker: HES1), glioblast (marker: GFAP), and neuroblasts (markers: NEUROD6) showed high expressions as indicated by Braun *et al*.^51^ (Supplementary Figure 1).

Aldinger *et al.* (2021) and Sepp *et al.* (2024) human single-cell cerebellar transcriptomes: Single-cell transcriptomes from the developing human cerebellum from Aldinger *et al.* and Sepp *et al.*^11,18^ were downloaded and combined. The datasets were checked to confirm that low quality cells had already been filtered out as published. Additionally, all 80,581 postnatal cells were filtered out from Sepp *et al.*, leaving a total of 169,549 cells across both datasets. Cell type annotations from the two datasets were consolidated and Seurat’s Canonical Correlation Analysis (CCA)^19^ was used to integrate the two datasets. Dimensionality reduction (principal component analysis (PCA) and uniform manifold approximation and projection (UMAP)) was performed on the combined cells, and clustered using Louvain clustering. Multiple clustering resolutions were explored and an optimal clustering resolution of 0.8 was selected for being the lowest resolution that identified the WLS+ KI67+ rhombic lip cluster. Clusters were then annotated based on expression of known markers and the original cell type annotations from the two datasets (Supplementary Figure 2a-c, Supplementary Table 1).

Nowakowski *et al.* (2017) human developing brain single-cell transcriptomes: Processed expression data for 4,262 developing brain cells^20^ were downloaded from the UCSC Cell Browser (https://cells.ucsc.edu/?ds=cortex-dev&cell=S41.F4). UMAP coordinates were downloaded from https://cells.ucsc.edu/cortex-dev/UMAP.coords.tsv.gz. Analysis was only performed on cells from RegionName “Cortex” (3,062 cells), and the “WGCNACluster” attribute was used to assign cell types. When possible, cell types were grouped together into broader classes, including: newborn excitatory neurons (nEN), newborn inhibitory neurons (nIN), mature excitatory neurons (EN), mature inhibitory neurons (IN), and medial ganglionic eminence (MGE) cells. Astrocytes, endothelial, microglia, OPC, and mural cells were grouped together as “non-neuronal”. Radial glial cells are enriched for HES1 and NES, dividing intermediate progenitor cells for EOMES (TBR2), and excitatory neurons for NEUROD6 (Supplementary Figure 2d).

Li *et al.* (2018) bulk RNA-seq of human brain through lifespan (“PsychENCODE, BrainSpan 2.0”): Processed mRNA-seq data from Li *et al.*^21^ was downloaded from the PsychENCODE project website and processed as described in our previous work^22^. For the grouped age analysis, groupings were constructed as follows: samples at or earlier than 12 post-conception weeks (PCW) were labelled as “first trimester”; those between 12 and 23 PCW, as “second trimester”; those between 24 and 37 PCW, as “third trimester”; samples 0-13 postnatal years were labelled “0-12Y”; those between 13 and 21 years were labelled, “13-20Y”, and those older than 21 years were labelled “>21Y”.

### Gene regulatory network analysis

pySCENIC^23^ (v0.12.0, Python v3.10.6) was used to infer modules of correlated gene expression in RL lineage cell clusters in the developing hindbrain dataset^11^ and in medulloblastoma^1^. A custom Enrichment Map gene set was used (downloaded from https://download.baderlab.org/EM_Genesets/January_04_2025/Human/symbol/Human_GOBP_AllPathways_noPFOCR_no_GO_iea_January_04_2025_symbol.gmt)^24^) which included pathways from GO Biological Processes (excluding those inferred from electronic annotation (IEA))^25^, HumanCyc^26^, MSigDB (hallmark, C2, and C3 collections)^27^, NCI-Nature^28^, NetPath^29^, Panther^30^, Pathbank^31^, Reactome^32,33^, and WikiPathways^34^. Only pathways with 10-250 genes were included (8,433 pathways). Network visualization was performed using Cytoscape^35^ v3.10.2 with EnrichmentMap v3.5.0 and AutoAnnotate v1.3.5.

### Medulloblastoma scRNA-seq analysis

Medulloblastoma scRNA-seq count expression matrices from Vladoiu *et al.* were provided by Dr. Michael Taylor. The dataset contained 27,735 cells from eight MB tumours (two SHH, two G3, and four G4). The count expression matrices from all cells were combined and normalized using SCTransform^36^. Batch correction was performed using fastMNN^37^ and the top 50 MNN-corrected principal components were used for UMAP dimensionality reduction and projection. Seurat’s FindMarkers was used to identify differentially expressed genes in selected clusters after adjusting for patient sex (logfc.threshold=0, min.pct=0.01, assay=”SCT”, recorrect_umi=FALSE, method=“LR” latent.vars=“Sex”). Seurat’s AddModuleScore was used to compute the score for CancerSea^38^ pathways for each cell, and their means were calculated for each cluster. PySCENIC^23^ was run to identify gene regulatory networks associated with each tumour cell cluster (see “Gene regulatory network analysis”).

### Medulloblastoma bulk tumour mutation analysis

Simple somatic mutations for medulloblastoma tumours were downloaded from the ICGC data portal^39^ on 5 October 2023 (https://dcc.icgc.org/releases/current/Projects/PBCA-DE, https://dcc.icgc.org/releases/current/Projects/PEME-CA). Gencode v19 (GRCh37) was used to ascertain exon and gene coordinates for genes of interest. Gene extent was defined from transcription start site to transcription end site. Molecular subgroup information for these tumour samples was downloaded from Northcott *et al.*^40^. The SpliceAI plugin from the Ensembl Variant Effect Predictor (VEP) was used to predict the impact of mutations on splice sites^41,42^.

### Medulloblastoma transcriptomics analysis

Normalized expression profiles from 768 primary Group 3 (G3) and Group 4 (G4) medulloblastoma (MB) tumours^43^ were downloaded from GEO (GSE85217) and tumour ploidy calls were obtained from Dr. Michael D. Taylor^43^. Sample metadata and somatic arm-level copy number aberration (CNA) information were obtained from supplementary tables associated with the same publication. A somatic chromosome i17q event was defined by the co-occurence of p-arm deletion and q-arm gain in the same tumour sample. Samples with missing chromosome 17 somatic arm-level CNA, gender, or age information were excluded, resulting in 714 tumours that were used in this analysis (64 WNT, 210 SHH, 133 G3, and 307 G4). A generalized linear model was used to test associations of BRCA1 expression with MB tumour subtypes, age, and sex. Survival analysis was limited to G3 and G4 MB tumours. After further excluding samples with missing survival data, death status, or other covariates, 382 tumours (111 G3 and 271 G4) were included in survival analysis. A multivariate Cox regression was performed, including overall survival in months and mortality status as the independent variables, and normalized BRCA1 expression level, age and sex as explanatory variables. We pooled both G3 and G4 tumours together for the analysis and included their individual subgroups as a covariate.

## Results

### Computational prediction of putative candidates for stalling differentiation

To study the stalled differentiation phenomena in the context of brain development and as a driver for Group 3 and 4 medulloblastoma (MB), we built a computational pipeline to identify transcription factors (TF) and epigenetic regulators that are upregulated in both the developmentally earlier rhombic lip (RL) progenitor cells relative to differentiating neurons, and in MB tumours (Figure 1a-b). For this we analyzed RL lineage cells from the developing human hindbrain^11^ (9 samples; 9-20 post-conception weeks, 9,208 cells). We identified 1,048 genes that were either upregulated in the RL subventricular zone cells relative to differentiating early unipolar brush cells (Seurat^19^, Q < 0.05; Supplementary Table 2), or that encoded transcription factors predicted to regulate these genes (AME^12^, Supplementary Table 3). After limiting genes to those encoding known epigenetic regulators or transcription factors^13,15^, and excluding DepMap essential genes^16^, 273 targets remained. We then rank-ordered targets by a proliferation-decreasing effect and average transcription level in the G3 MB cell line D425^16^, and selected genes until all putative medulloblastoma driver genes were included. The list of putative drivers includes those previously identified as being recurrently mutated or aberrantly overexpressed. This pipeline resulted in a total of 252 target genes that we deem potential candidates for driving stalled differentiation in Group 3 and 4 medulloblastoma (Figure 1c, Supplementary Table 4). The top candidate on the list was OTX2, a developmental TF that defines midbrain identity in early gestation, that is well known for being overexpressed in medulloblastoma tumours, and the knockdown of which results in differentiation of Group 3 medulloblastoma cell lines^2,44^. Surprisingly, we observed that high-ranked candidate genes included BRCA1, best known for its role in DNA damage repair and loss-of-function mutations driving risk of familial breast and ovarian cancer^45–47^. On further investigation, we observed that BRCA1 expression appeared localized to dividing rhombic lip progenitors (Figure 1d-e). BRCA1 has been previously reported to control neurogenesis in the developing mouse cerebral cortex, hippocampus and cerebellum^48–50^, but its role in the developing human brain remains unexplored. This gap in the role of BRCA1 in human brain development prompted us to investigate BRCA1 more closely in this context.

**Figure 1.**
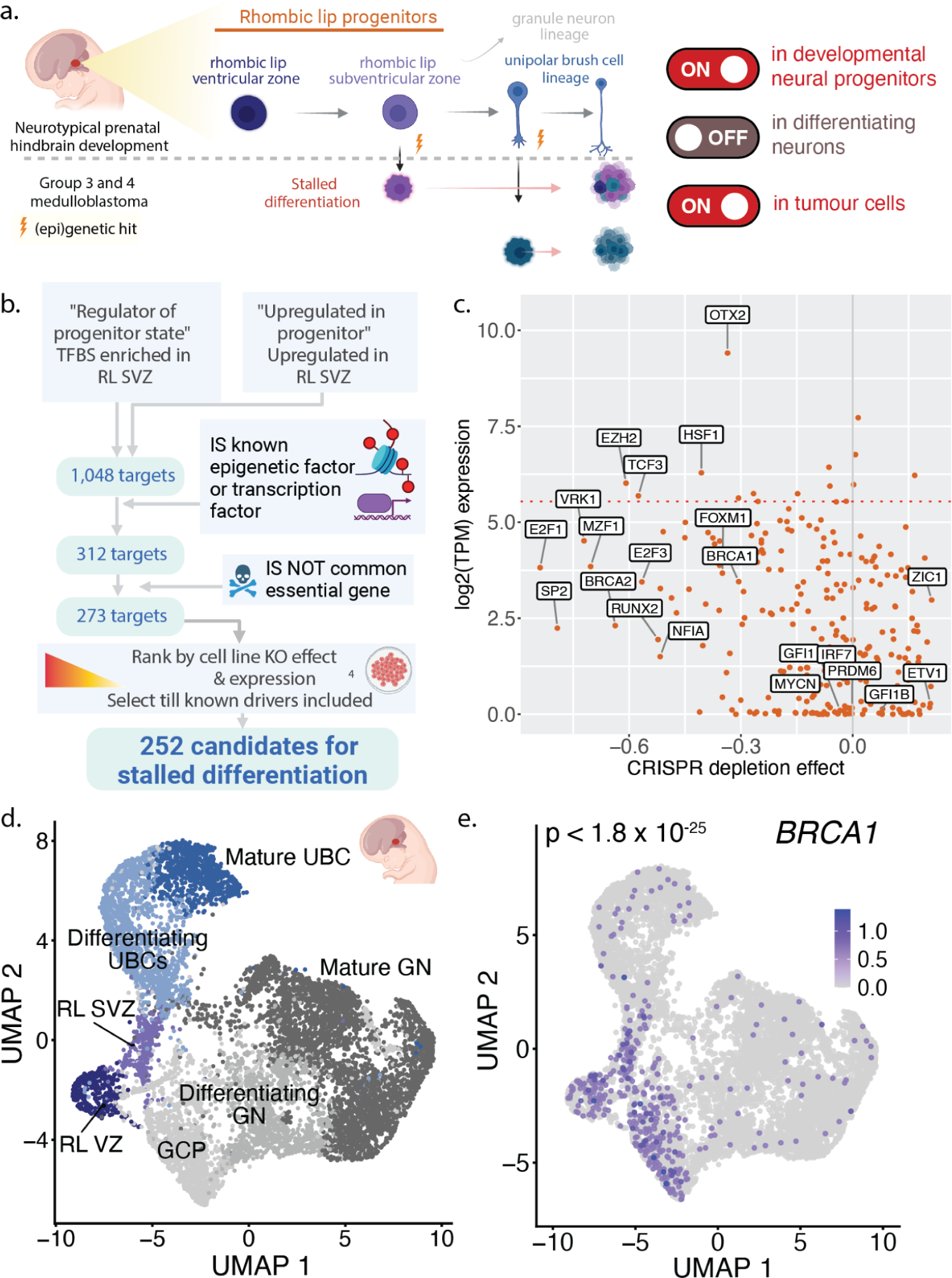
Integration of human developing cerebellum and medulloblastoma single-cell transcriptomes to nominate candidates promoting stalled differentiation states in cancer. a. Stalled differentiation model for tumour initiation in Group 3 and 4 medulloblastoma, and expected pattern of transcription of a mediator of stalled differentiation. b. Computational pipeline to identify putative candidates for stalled differentiation. c. Expression and CRISPR depletion effect of the top 75 targets in the D425 Group 3 medulloblastoma cell lines (DepMap); highlighted genes are known medulloblastoma drivers. d. UMAP of rhombic lip lineage cells from the developing human hindbrain (25,213 cells). e. *BRCA1* expression in human rhombic lip cells. *p*-value from differential expression of *BRCA1* in rhombic lip subventricular cells versus differentiating UBCs. RL VZ: rhombic lip ventricular zone; RL SVZ: rhombic lip subventricular zone; GCP: granule cell progenitors: GN: granule neurons; UBC: unipolar brush cells.

### *BRCA1* is enriched in neural stem and progenitor cells in early human neurodevelopment

To ascertain how broadly *BRCA1* is expressed in the human brain, we mapped *BRCA1* expression at cellular resolution from four independent single-cell transcriptomic datasets in human brain development for a total of 142 brain samples, 9 regions, and 1.77 million cells, including in a multi-locus transcriptomic dataset that spans the human lifespan (Figure 2a, sample sizes for all cell clusters and tissue samples used for statistical tests in Supplementary Table 5)^18,21,51,52^. We found that in the developing brain, *BRCA1* expression is consistently enriched in dividing neural stem and progenitor cells (Figure 2b-d). In the developing cortex, *BRCA1* is significantly upregulated in radial glial cells and intermediate progenitor cells, as compared to newborn excitatory or inhibitory neurons (Figure 2b, p < 2×10^−16^, one-tailed t-test, mean of 598 cells per cluster, range [75-1,532]). This pattern is also true in the developing hindbrain, where *BRCA1* expression is significantly higher in the rhombic lip neurogenic niche, as compared to the differentiating neuronal lineages it gives rise to, the unipolar brush cells and granule cell lineage (Fig 2c, p < 2×10^−16^, one-tailed t-test, mean of 8,612 cells per cluster [1,783-17,205]). For this hindbrain analysis, we integrated cells from a second, independent dataset^18^ to the developmental dataset previously used^11^. The effect does not appear to be sex-specific, as rhombic lip cells from both male and female samples show *BRCA1* enrichment (p < 0.001, two-tailed WMW test, Supplementary Figure 3).

**Figure 2.**
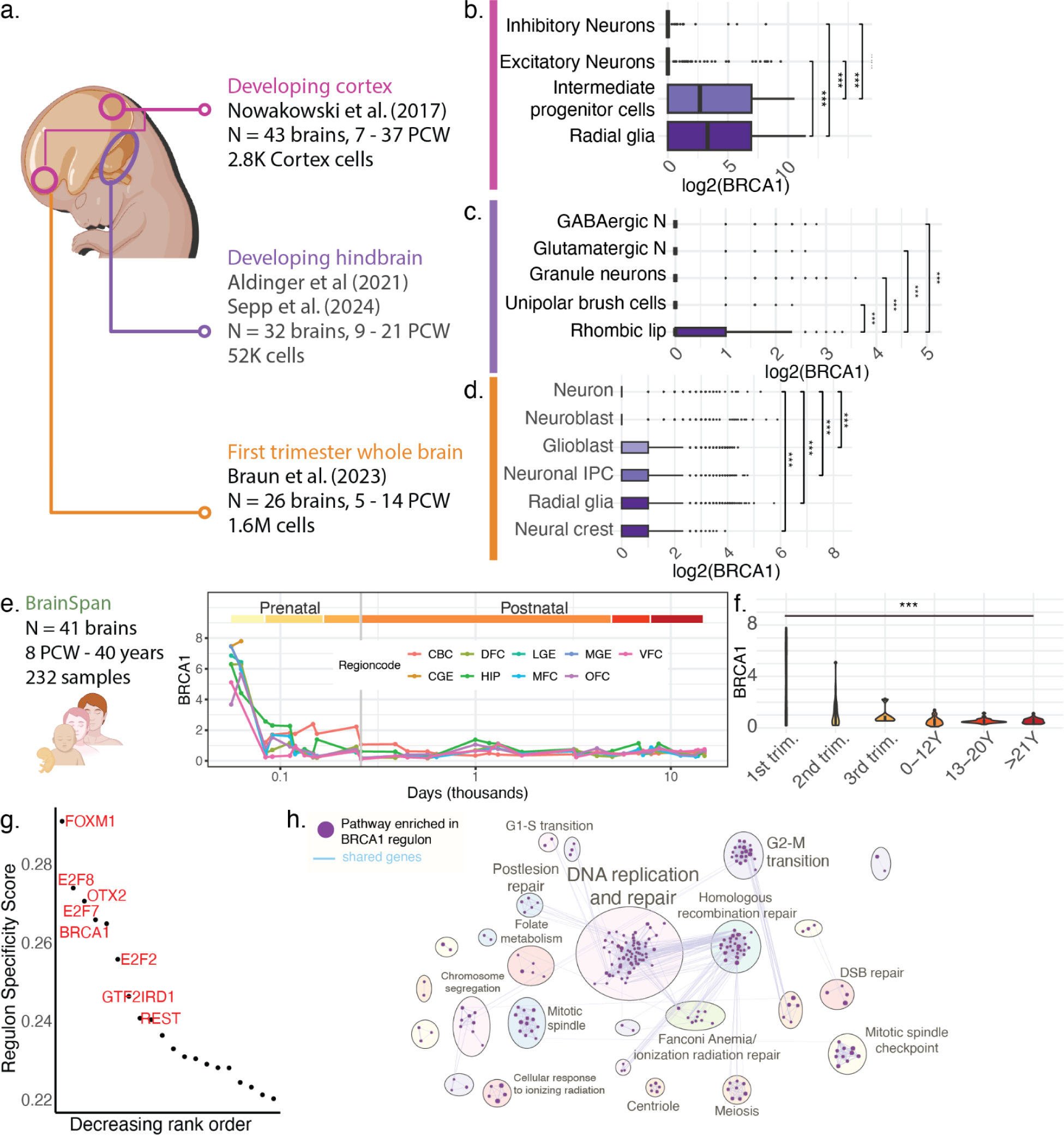
BRCA1 expression is enriched in dividing neural cells in the early prenatal human brain and is correlated with cell cycle and DNA damage repair processes. a. Developing brain single-cell transcriptome datasets analyzed. b. Normalized expression of BRCA1 by major cell clusters in the developing cortex c. the developing hindbrain, and d. the entire brain in the first trimester. Boxplots show normalized expression and p-values are from a one-tailed t-test. e. Normalized expression of BRCA1 through the human life span. Lines show data for different brain regions, and coloured rectangles indicate different age ranges. f. BRCA1 expression in Brainspan dataset grouped by ages. p-value from one-way ANOVA. g. Gene regulatory networks driving rhombic lip subventricular zone in the developing human hindbrain^11^. h. Pathways enriched in genes comprising the BRCA1 regulon in the human rhombic lip subventricular zone (g:Profiler, Q < 0.05). Nodes are pathways, edges indicate shared genes. Labels highlight major themes in pathway clusters. Sample sizes for panels b-d,f in Supplementary Table 5. PCW: post-conception weeks. IPC: intermediate progenitor cells; Glutam. Neuron: glutamatergic neuron; GABA neuron: GABAergic neuron; CBC: cerebellar cortex; DFC: dorsal frontal cortex; LGE: lateral ganglionic eminence; MGE: medial ganglionic eminence; VFC: ventral frontal cortex; CGE: central ganglionic eminence; HIP: hippocampus; MFC: medial frontal cortex; OFC: orbitofrontal cortex.

In the first trimester, *BRCA1* expression is significantly higher in dividing neural progenitors, including neural crest cells, radial glia, neuronal intermediate progenitor cells, and glioblasts, as compared to neuroblast or neurons (Figure 2d, all comparisons, 2×10^−16^, one-tailed t-test). Across the human lifespan and multiple brain regions, *BRCA1* expression is the highest in the first trimester of prenatal development compared to later in development or in postnatal life (Figure 2e-f, p < 2×10^−16^, one-way ANOVA), a trend that appears to apply to the cortex, cerebellum, and hippocampus, among other regions of the brain (Figure 2e).

To ascertain which cellular processes BRCA1 may mediate in development, we identified gene regulatory networks driving the rhombic lip subventricular zone, where BRCA1 expression is enriched. BRCA1 itself was identified as among the top regulons of the rhombic lip subventricular zone (Figure 2g, Supplementary Tables 6-7). Notably, the subnetwork of genes correlated with *BRCA1* expression were enriched in processes of cell proliferation, the mitotic cell cycle, DNA replication, homologous recombination repair, and chromosome segregation (Figure 2h; g:Profiler, Q < 0.05, Supplementary Table 8).

### In medulloblastoma, *BRCA1* expression increase is associated with i17q status, Group 4 tumours, and cell cycle activity

To understand the nature of BRCA1 misregulation in medulloblastoma, we next examined the genetic and transcriptomic variation of this gene using the largest published survey of medulloblastoma to date (Fig. 3a, N=714 tumours)^40,43^. We observed gene body mutations of *BRCA1* in 1.5% of Group 3 MB and 3% of Group 4 MB tumours^40^ (N=2/131 and 6/193 tumours, respectively); none of these mutations were located in exons (Supplementary Figure 4) and none are predicted to affect splice sites (Supplementary Table 16). The *BRCA1* gene is located on chr17q, and isochromosome 17q (i17q) aberration is observed in ∼27% of Group 3 MB tumours and 57% of Group 4 MB tumours^43^. We therefore investigated if *BRCA1* expression was increased in Group 3 and 4 medulloblastoma tumours carrying the i17q mutation status versus those that did not. We observed that *BRCA1* is indeed significantly increased in i17q carriers (Figure 3b; p < 7.7×10^−11^; total 440 tumours, 214 carriers; GLM adjusting for age, sex and subgroup), and that this effect is seen in tumours from male and female patients (p < 7.8×10^−5^, Supplementary Figure 5a). The increase in i17q carriers is observed even after controlling for tumour ploidy (N=319 tumours with ploidy information; p < 7.6×10^−9^ for i17q status, Supplementary Figure 5b), and separately, when tumours are separated into Group 3 and 4 subgroups (N=133 Group 3 tumours, p < 2.1×10^−5^ and N=307 Group 4 tumours, p < 4.6 x 10^−7^, respectively; Supplementary Figure 5c).

**Figure 3.**
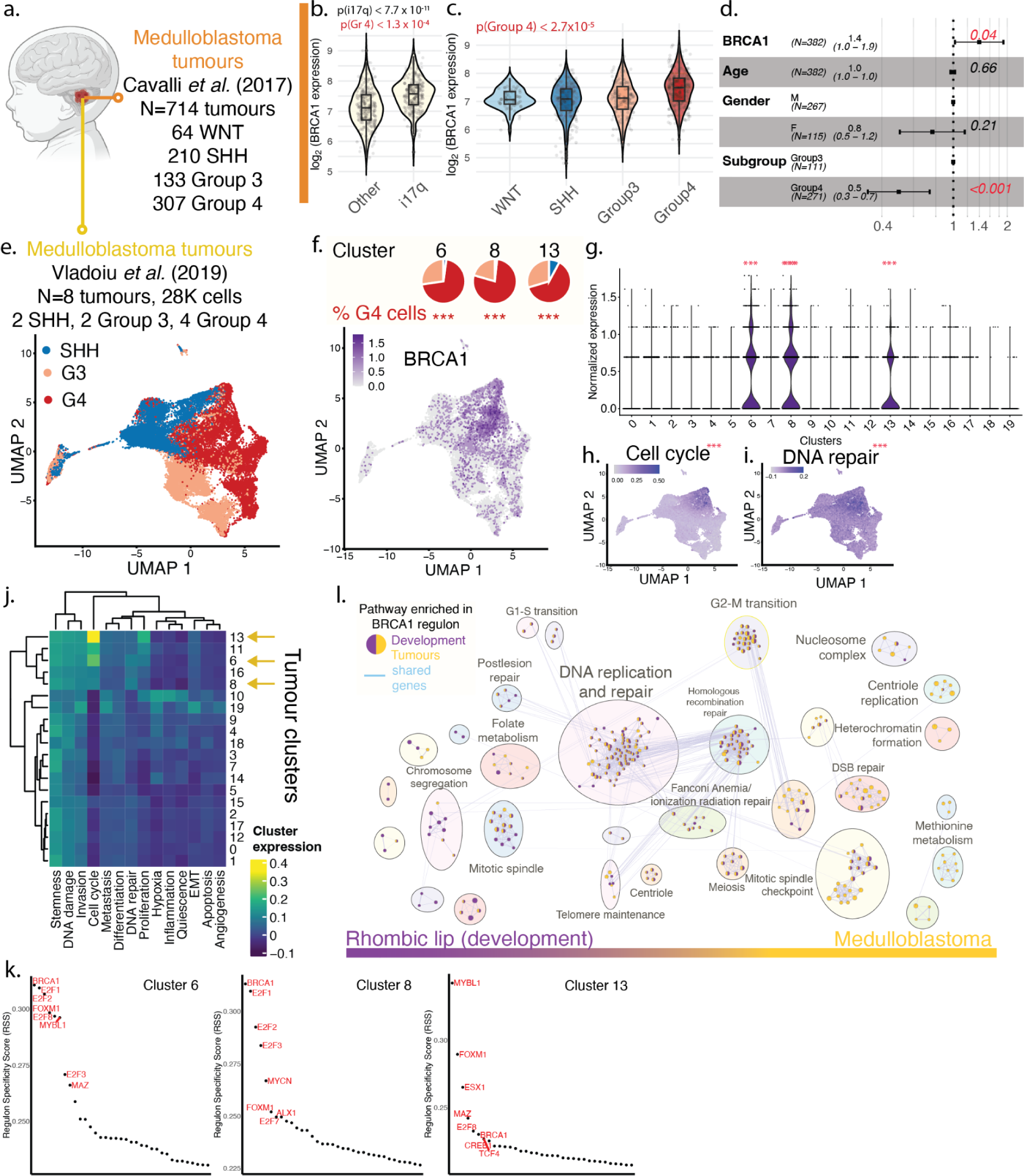
BRCA1 expression in medulloblastoma tumour genomes. a. Dataset used for the analysis (in b, c, d). b. BRCA1 expression in Group 3 and 4 medulloblastoma tumours for carriers of isochromosome 17q (N=214 tumours) versus non-carriers (N=226 tumours); marginal p-values from GLM model after adjusting for age, sex, and subgroup. c. BRCA1 transcription in bulk transcriptomes of medulloblastoma tumours divided by its 4 subtypes. p-value for effect size in Group 4 tumours (GLM model after adjusting for age, sex, and subgroup). d. Hazard ratio for overall survival in Group 3 and 4 medulloblastoma tumours (N=382 tumours). e. Medulloblastoma single-cell transcriptomes dataset details. f. Expression of BRCA1 in tumour clusters, and proportion of cells of each subtype in the three tumour clusters demonstrating significant upregulation of BRCA1. Pie charts show cellular composition of these clusters and asterisks indicate statistical significance for enrichment of Group 4 medulloblastoma cells in each cluster (p < 0.001, Fisher’s Exact Test). g. Normalized expression of BRCA1 in tumour clusters, asterisks indicating significance in differential expression of BRCA1 in each cluster (p < 0.001, FindMarkers). h-i. Expression of CancerSea modules for (h) Cell cycle (137 genes) and (i) DNA repair (119 genes) in tumour cells. Asterisks indicate statistical significance of correlation of module score and BRCA1 expression (*t*-test). j. Correlation of medulloblastoma tumour clusters with CancerSEA modules. Clusters showing upregulation of BRCA1 are highlighted with arrows. k. Top transcription factors predicted to drive gene expression in BRCA1-upregulated medulloblastoma cells (PySCENIC). l. Pathways enriched in BRCA1 regulon in development and medulloblastoma. Nodes show enriched pathways and edge connect pathways with shared genes, and nodes are colour-coded based on whether pathways were detected in development (purple), cancer (orange), or both (split-coloured circle) (gProfiler, Q < 0.01 shown).

However, even after adjusting for i17q status, a significant association with Group 4 medulloblastoma subgroup remains (marginal p for tumour subgroup, p < 1.3×10^−4^); this association also remains after adjusting for tumour ploidy (p < 2.3×10^−3^ for Group 4 subgroup). Consistent with this observation, *BRCA1* expression is increased in Group 4 medulloblastoma tumours, relative to WNT, SHH, and Group 3 medulloblastoma tumours (Figure 3c, p < 2×10^−5^ effect for tumour subgroup; generalized linear model (GLM) adjusted for age, sex and tumour subgroup; N=714 tumours; 64 WNT, 210 SHH, 133 Group 3, 307 Group 4). The increase in *BRCA1* expression is observed in Group 4 tumours from male and female patients (p < 2.1×10^−3^, two-tailed Wilcoxon-Mann-Whitney (WMW) test; Supplementary Figure 5d). Within tumour subgroups, at the tumour subtype level, an increase in BRCA1 expression is significantly associated with Group 4α or Group 4β subtype (Supplementary Figure 5e, p < 5.0×10^−9^ and p < 5.3×10^−4^, GLM; age, sex, tumour subtype adjusted; 19-109 tumours per subtype; mean=59.5); no association with Group 4γ was observed (p > 0.1).

The association between *BRCA1* expression and overall survival is less clear. When Group 3 and 4 medulloblastoma tumours are examined together, increased BRCA1 expression is associated with poor prognosis (Figure 3d; HR=1.40; 439 tumours, adjusting for age, sex, and subgroup). However, this effect is lost when considering only Group 3 or only Group 4 tumours (p>0.1, Supplementary Figure 5f-g).

We next examined *BRCA1* expression in medulloblastoma single-cell transcriptomes^1^ (Figure 3e-l, N=8 tumours, 27,735 cells). Three tumour cell clusters significantly overexpressed *BRCA1*, relative to other tumour clusters (Figure 3e-g, p < 2×10^−16^ for each cluster, adjusted for sex; Supplementary Figure 6, Supplementary Table 9-11). These clusters showed an enrichment for Group 4 medulloblastoma cells over other subtypes (p < 10^−32^, all three clusters; Fisher’s Exact Test; Supplementary Table 12). Of all the hallmark cancer attributes, these clusters are most correlated with expression of pathways involving cell division, including cell cycle and proliferation (Figure 3g-j); *BRCA1* expression is most strongly associated with expression of genes involved in DNA repair and cell cycle regulation (Figure 3h-i; Pearson’s rho=0.39 and 0.38 respectively, p < 2×10^−16^; t-test; Supplementary Figure 7). In these clusters, *BRCA1* is predicted to be a top transcription factor (Figure 3k, pySCENIC, Supplementary Tables 13-14). Genes co-expressed with *BRCA1* in these clusters are enriched for DNA replication and repair, including homologous recombination, DNA double stranded break repair, and the cell cycle checkpoint (g:Profiler, Q < 0.01; Figure 3l; Supplementary Table 15). Overall, *BRCA1* enrichment is observed in Group 4 medulloblastoma tumour cells at the bulk and single-cell transcriptomic level, and this enrichment is strongly associated with cell cycle and DNA damage repair activity.

## Discussion

BRCA1 is best known for its role as a tumour suppressor involved in DNA double-stranded break repair^10,53–55^. In a dividing cell, double-stranded DNA breaks can trigger cell cycle arrest and DNA repair through a network of proteins that involve break detectors such as p53 and ATM, repair mediators such as BRCA1, and G2/M arrest through interactions of p21 with cyclin dependent kinase 2. Brain development faces the challenge of maintaining genome integrity during extensive neural stem and progenitor cell proliferation; an extreme example of this challenge is the rhombic lip neurogenic niche in the hindbrain, which ultimately generates tens of billions of granule neurons in the adult brain^56,57^. Brain development is also accompanied by the production of reactive oxygen species (ROS), which induces DNA damage^58^. In the mouse, a *Brca1* knockout in neural stem cells results in a smaller brain size due to a reduced number of neurons in multiple regions of the brain, including the cerebral cortex, hippocampus, and cerebellum^48,49^. This effect can be rescued by a double knockout of *Brca1* and *p53* or *ATM*, suggesting that BRCA1 mediates neurogenesis via the DNA damage repair pathway^48^. Separately, the BRCA1 DNA damage repair pathway can be regulated to increase double stranded breaks and induce neural stem cells to exit from a proliferative cycle, for example, by ROS-mediated signaling^50^. However, all mechanistic studies to date have been performed in the mouse, and the role of BRCA1 in human brain development, to our knowledge, remains unreported to date.

In humans, germline loss-of-function mutations are associated with an increased risk of familial breast and ovarian cancer ^45–47^, as a secondary, somatic mutation in the gene can result in a loss of DNA repair, loss of genome integrity, and oncogenesis^10^. We note that the model consistent with our observations is one of increased BRCA1 expression in tumour cells, which may serve to confer genomic protection as these cells proliferate, rather than a loss-of-function scenario which would increase the risk of oncogenic mutations. Indeed, we do not observe exonic BRCA1 mutations in medulloblastoma genomes. Separately, medulloblastoma tumours are enriched in loss-of-function mutations in genes mediating DNA damage repair, including members of the Fanconi anemia pathway for Group 3 and 4 medulloblastoma tumours^2^. Interestingly, germline carriers of *p53* mutations and *BRCA2*^−/−^ have an increased risk of developing a different molecular subtype of medulloblastoma, the Sonic Hedgehog (SHH) subtype^59,60^. We hypothesize that these loss-of-function mutations in the DNA damage repair pathway in Group 3 and 4 medulloblastoma reflect an unrelated mechanism of oncogenesis.

In sum, our data suggest that BRCA1, and likely the DNA damage repair pathway, may have a cell-protective role during proliferation in neurodevelopment and subsets of Group 3 and 4 medulloblastoma. Our findings to date are consistent with the observations in mouse neurodevelopment, wherein the BRCA1-associated DNA damage repair pathway protects dividing neural stem cells against DNA damage caused by reactive oxygen species and replication errors, and may be a controller of neural stem cell fate^50^. Implications for neurodevelopment and cognitive development of individuals with genetic haploinsufficiency in the DNA damage repair pathway, including carriers of *BRCA1* mutations, will need to be explored. Mice heterozygous for *Brca1* demonstrate learning and memory and motor deficits^61^; this effect was shown to be exacerbated by *in utero* ethanol exposure, which increases ROS production. The impact of *BRCA1* germline mutations on human brain development is, to our knowledge, not well documented. Two case studies have reported neural migration defects and cerebral hypoplasia in germline carriers of *BRCA1* mutations; in both instances, this defect was attributed to somatic loss of heterozygosity early in development^62,63^.

Separately, our data suggest that BRCA1 may be supporting Group 3 and 4 medulloblastoma growth. In our study, increased *BRCA1* expression is associated with a modest risk of poor prognosis in Group 3 and 4 medulloblastoma tumours as a whole (Fig 3d), but demonstrates a strong association with i17q carrier status. As the gene encoding BRCA1 is located on chr17q, we speculate that duplication of the chr17q arm may contribute to the increased expression of *BRCA1* in these tumours. However, there appears to be a separate, measurable effect of increased *BRCA1* expression in Group 4 medulloblastoma tumours. Targeting the BRCA1-associated DNA damage repair pathway could be an attractive therapeutic strategy for BRCA1-overexpressing tumours, but this avenue requires further exploration.

## Data and Code Availability

Software used for analysis in this manuscript will be made publicly available upon publication.

## Supplementary Figures

**Supplementary Figure 1.**
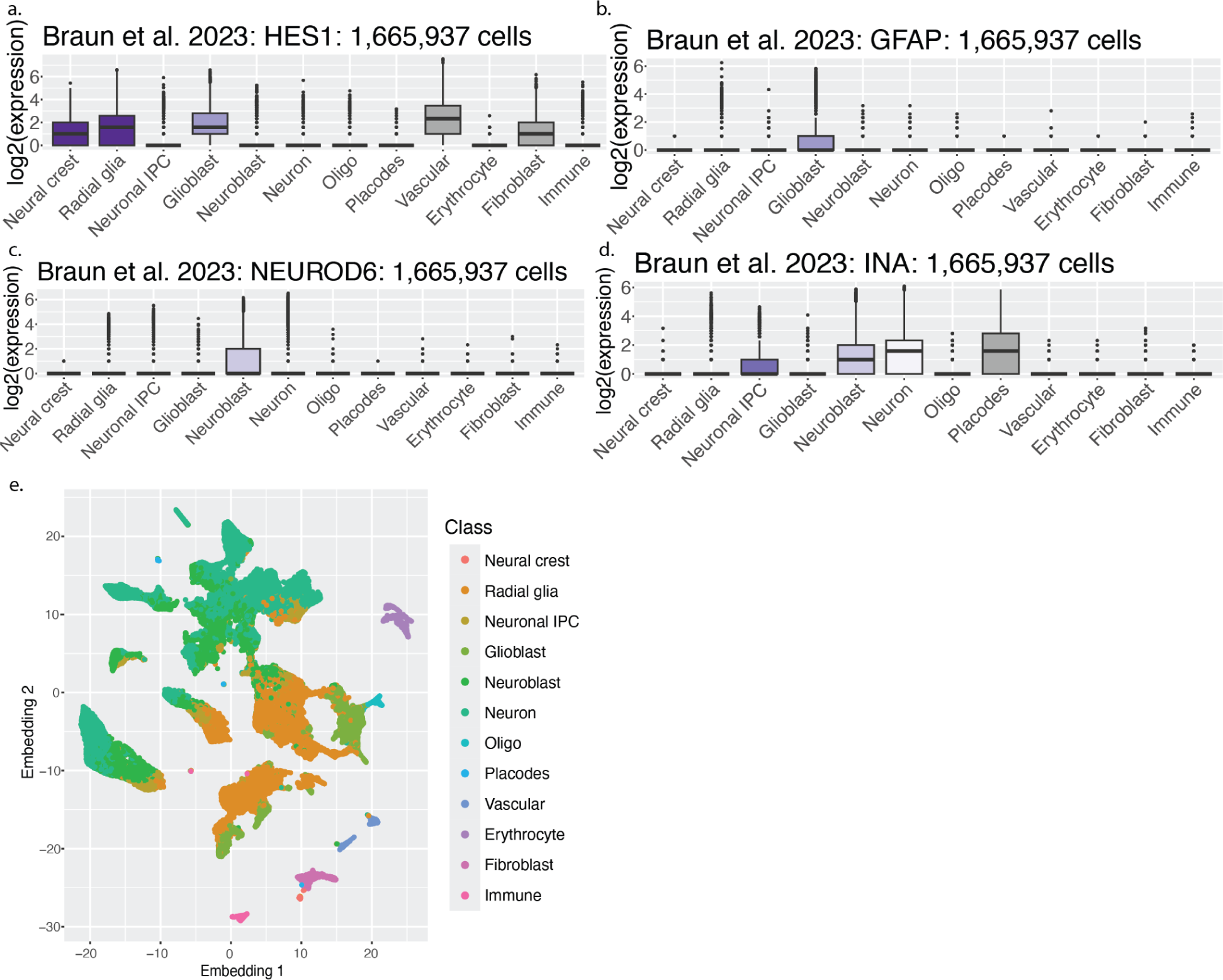
Expression of cell type markers and embedding for the Braun *et al.* 2023 first trimester dataset. Boxplots show cluster-wise expression of (a) *HES1*, a marker of radial glia; (b) *GFAP*, a marker of astrocytes, (c) *NEUROD6*, a marker of neuronal differentiation, and (d) *INA*, a marker of immature neurons. (e) Embedding of the 1.6 million cells from the Braun *et al.* dataset.

**Supplementary Figure 2.**
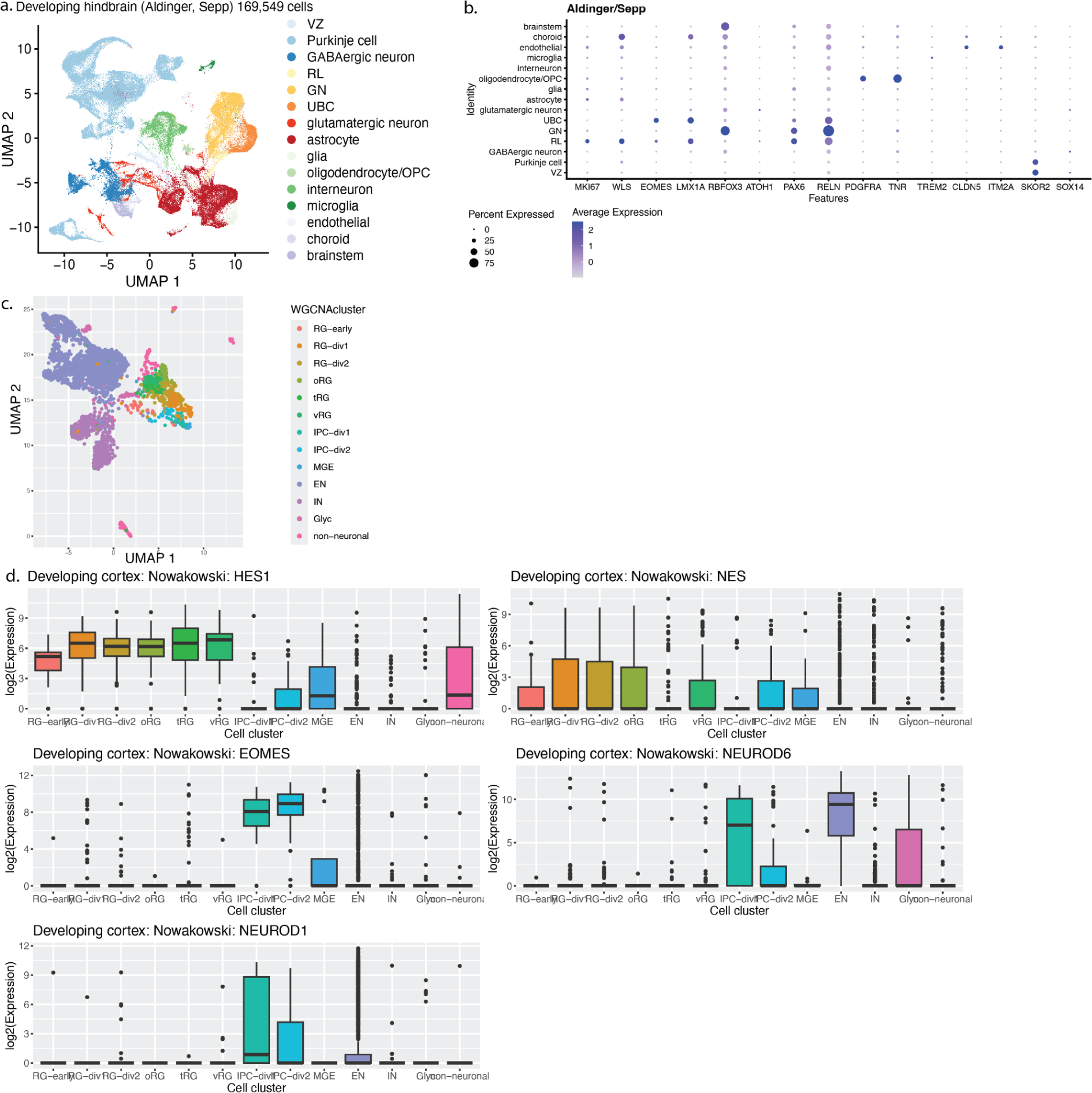
Clustering and cluster-specific markers for the human neurodevelopmental single-cell transcriptome sets used in this study. (a) Dimensionality reduction of the human developing cerebellum datasets from Aldinger *et al.* and Sepp *et al.* integrated by canonical correlation analysis (169,549 cells). RL: rhombic lip, VZ: ventricular zone, GN: granule neuron, UBC: unipolar brush cell. (b) Marker genes expressed in cell clusters of the integrated dataset. (c) Dimensionality reduction of the developing cerebral cortex dataset from Nowakowski. *et al.* (3,062 cells). (d) Marker genes expressed by different cell clusters. RG: radial glia; vRG: ventral radial glia; tRG: transverse radial glia; oRG: outer radial glia IPC: intermediate progenitor cell; MGE: medial ganglionic eminence; EN: excitatory neuron; IN: inhibitory neuron

**Supplementary Figure 3.**
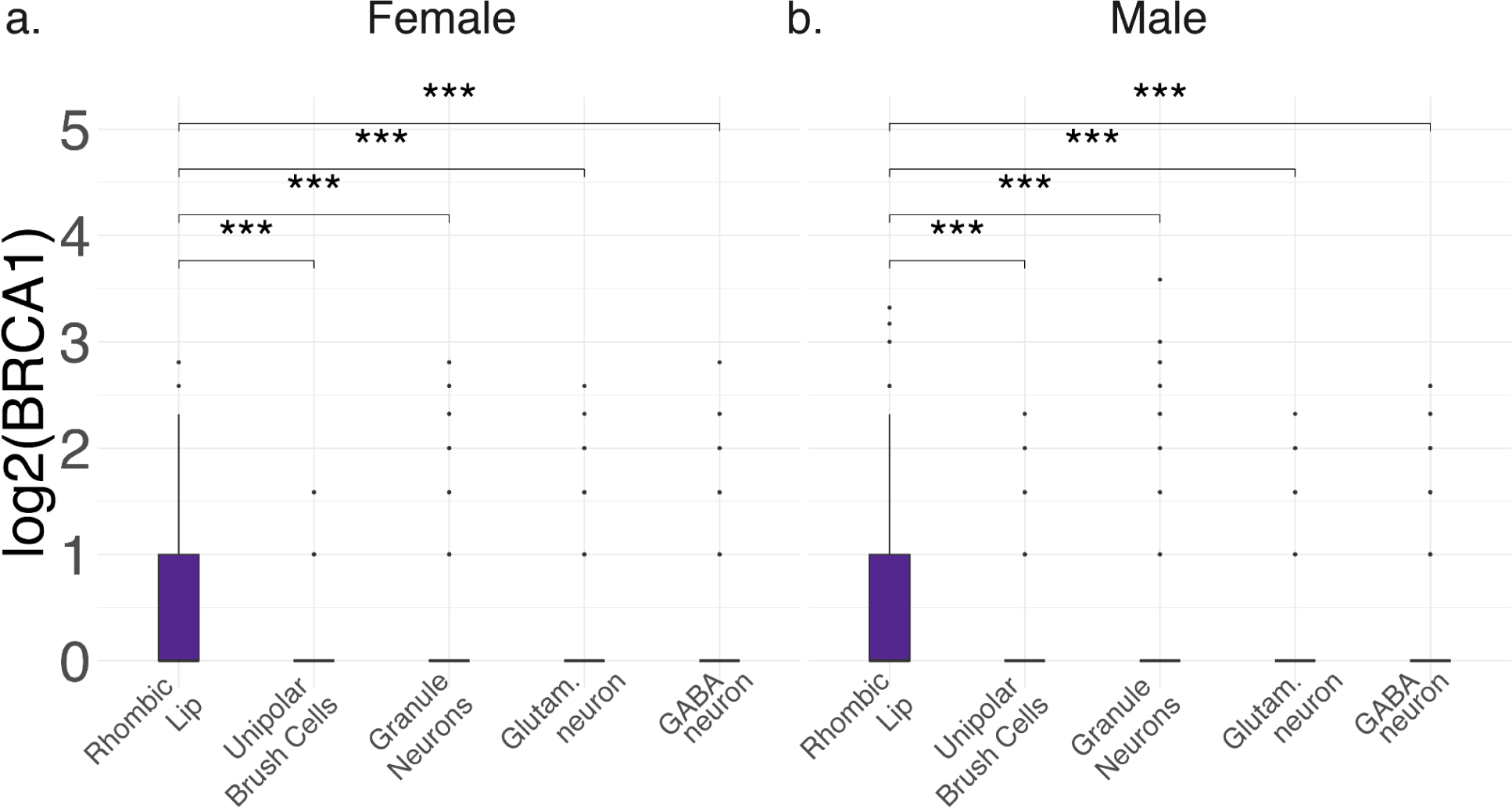
Sex-stratified normalized BRCA1 expression in human rhombic lip neural cell clusters. a. Cells from female samples (N=14 samples, total 22,273 cells); b. Cells from male samples (N=16 samples, total 26,133 cells). P-values from two-tailed WMW test. Sample sizes for groups shown in Supplementary Table 5. *** p < 0.001.

**Supplementary Figure 4.**
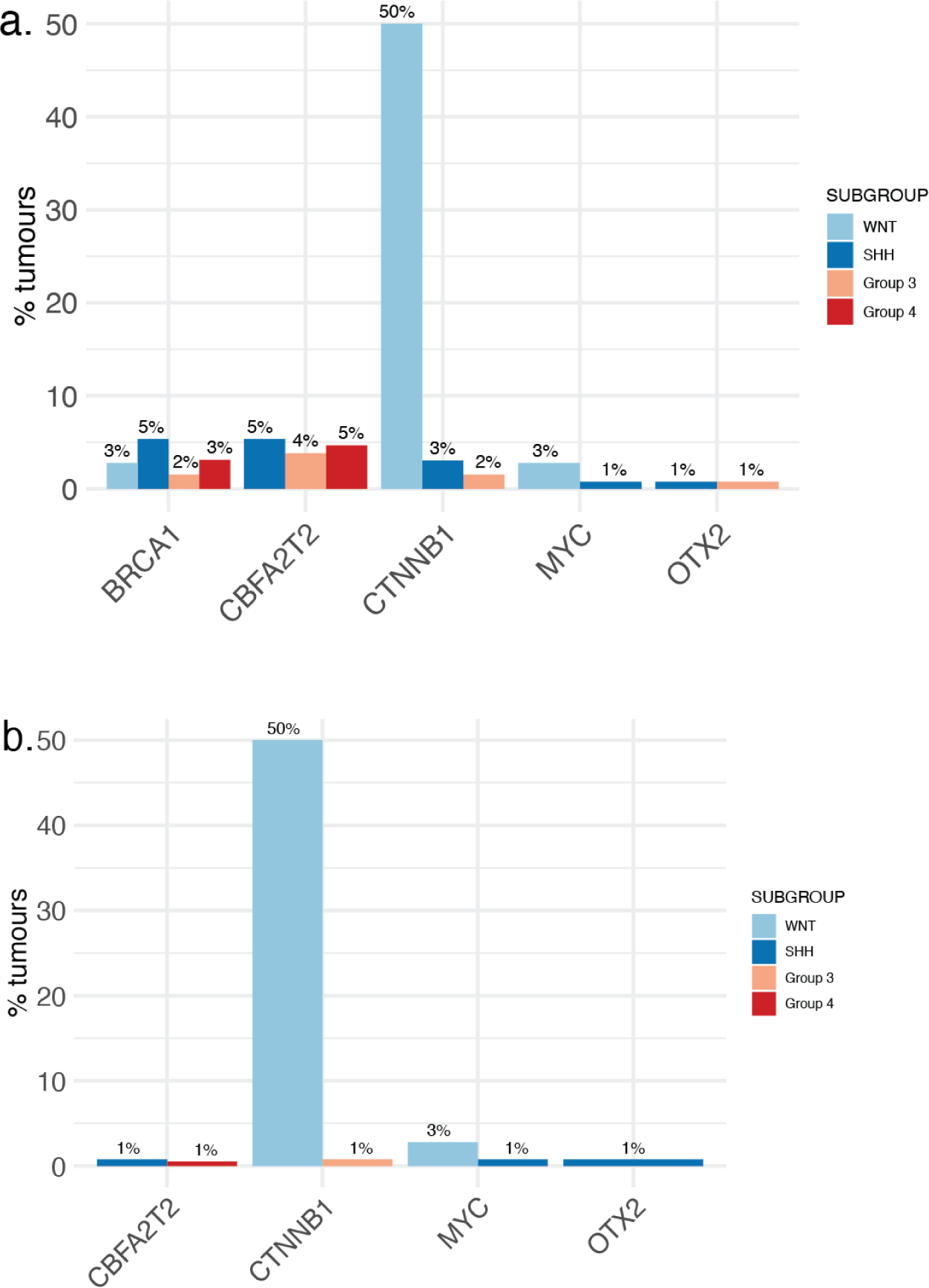
Percent tumours carrying a. gene body mutations and b. exonic mutations in *BRCA1* and selected medulloblastoma driver genes for comparison. Total 491 tumours; WNT=36 tumours; SHH= 131 tumours; Group 3 = 131 tumours; Group 4 = 193 tumours.

**Supplementary Figure 5.**
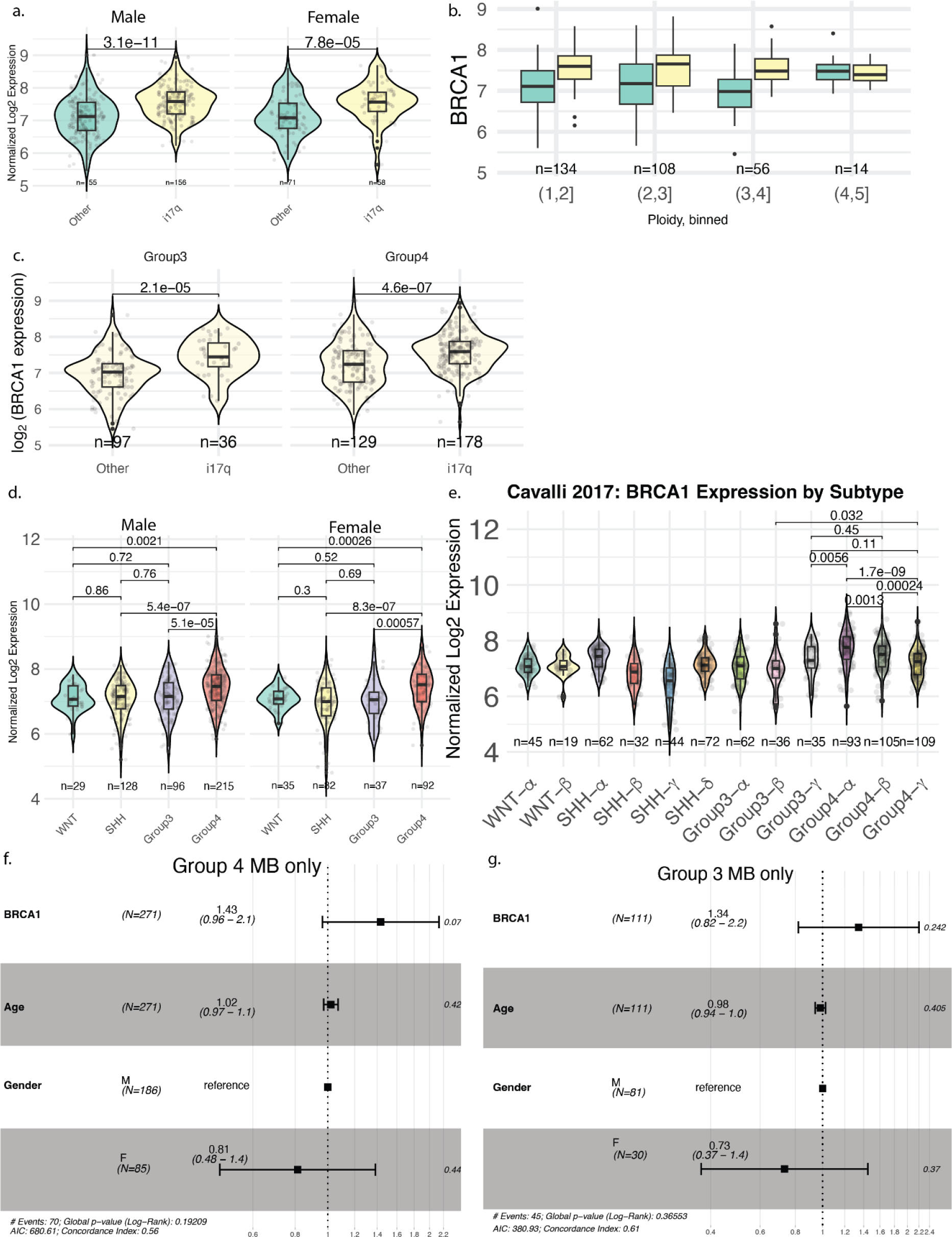
BRCA1 expression in bulk RNAseq of medulloblastoma tumours. a. BRCA1 expression in Group 3 and 4 MB tumours with i17q aberration (yellow) versus tumours without this chromosomal aberration (green), stratified by sex. For all panels in this plot, nominal p-values are from two-tailed WMW tests. Sample sizes for each group shown under corresponding violin plot. b. BRCA1 expression in Group 3 and 4 MB tumours by binned tumour ploidy, and stratified by i17q carrier status. c. BRCA1 expression in i17q carriers versus others, stratified by tumour subgroup. d. BRCA1 expression of tumours, stratified by tumour subgroup and sex. d. BRCA1 expression by tumour subtype and stratified by sex. e. Cox proportional hazards of BRCA1 expression in Group 4 MB tumours. f. Cox proportional hazards of BRCA1 expression in Group 3 MB tumours.

**Supplementary Figure 6.**
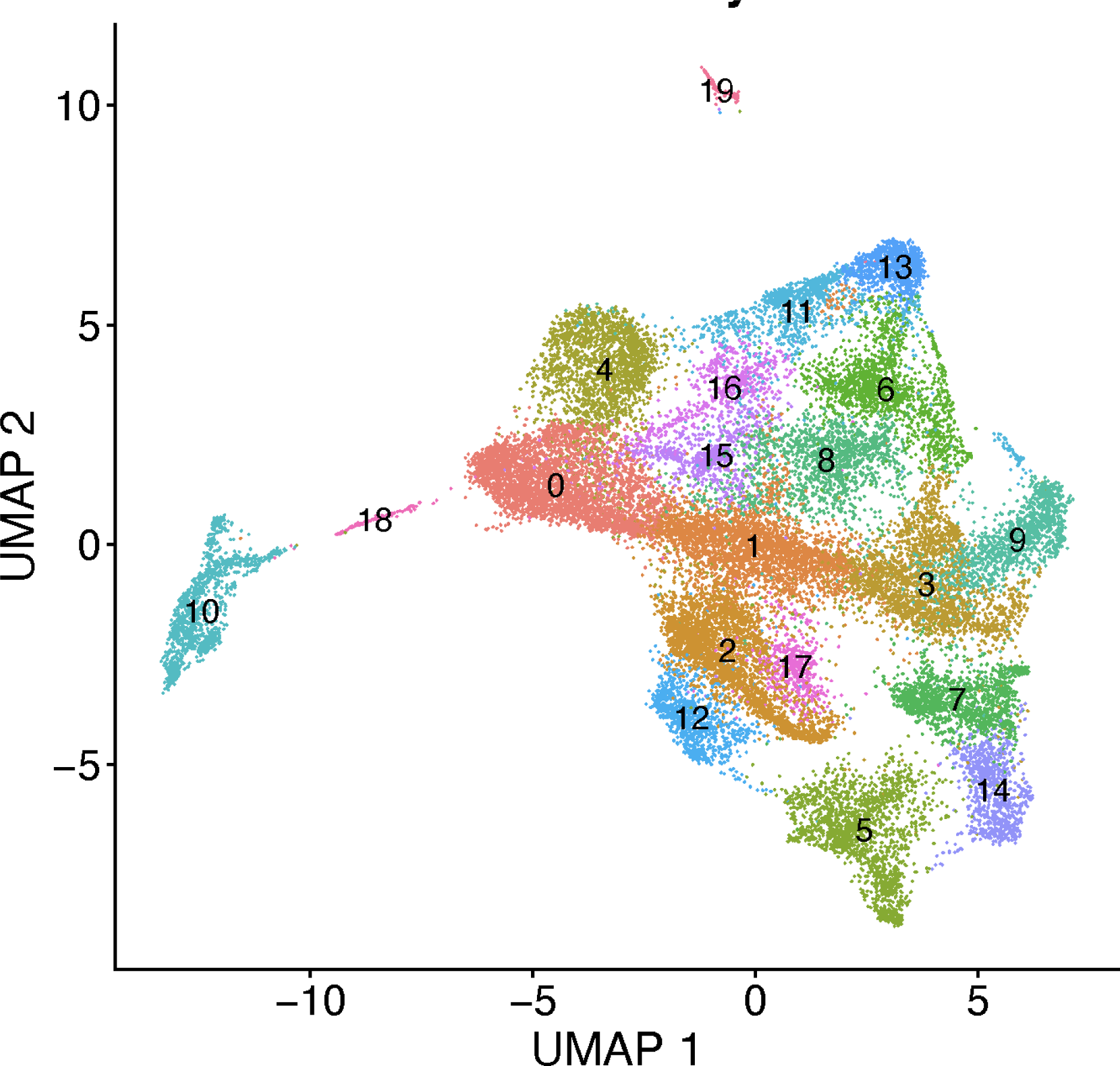
UMAP of tumour clusters for the Vladoiu et al.^1^ medulloblastoma scRNAseq dataset.

**Supplementary Figure 7.**
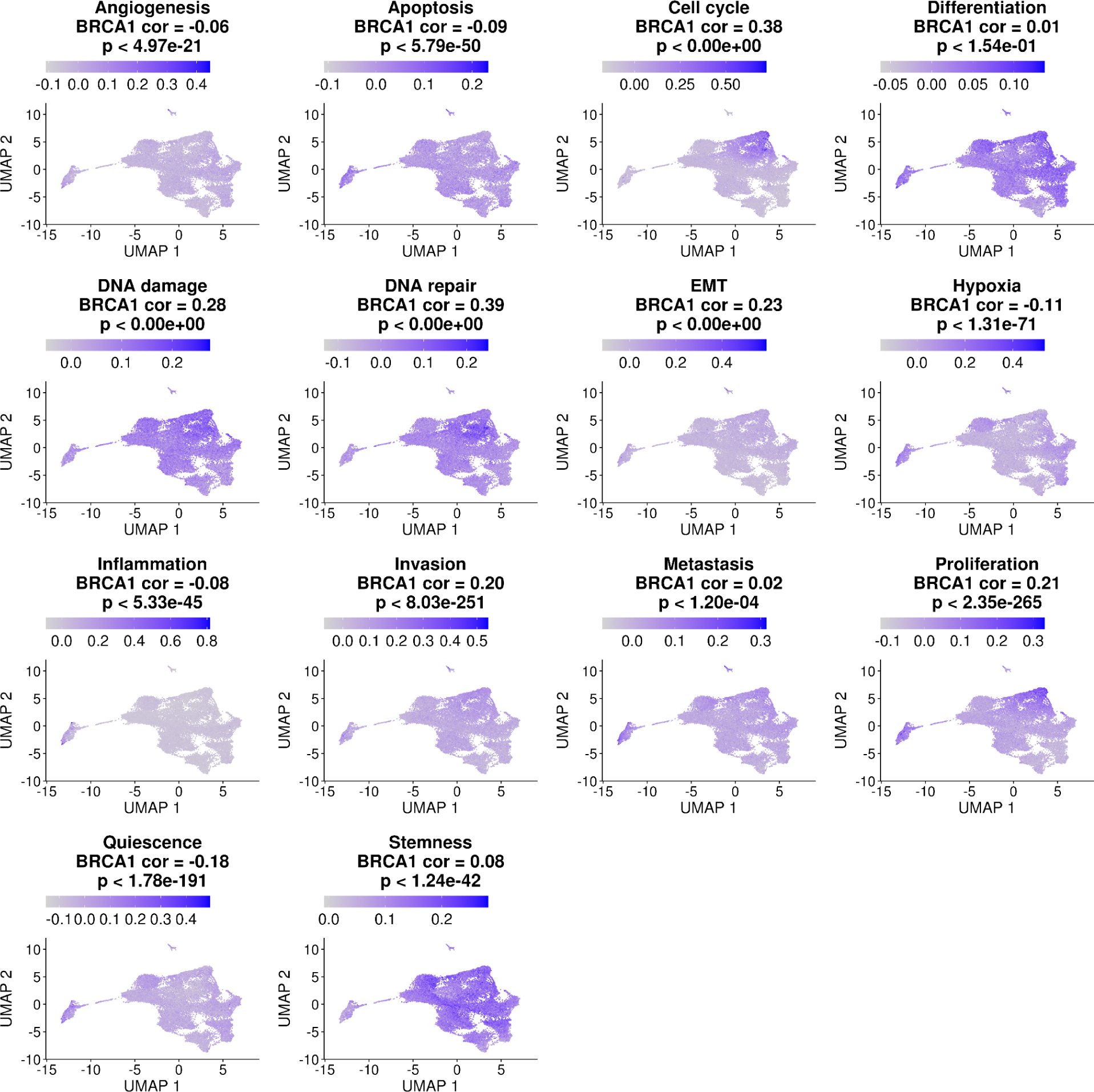
Correlation of BRCA1 expression in medulloblastoma tumour cells and expression of CancerSEA^38^ pathway modules. Each panel shows the relative expression level of a CancerSEA module. The title indicates the Pearson correlation coefficient of the expression of the corresponding module with BRCA1 expression, and associated statistical significance.

